# Predicting Individual Task Contrasts From Resting-state Functional Connectivity using a Surface-based Convolutional Network

**DOI:** 10.1101/2021.04.19.440523

**Authors:** Gia H. Ngo, Meenakshi Khosla, Keith Jamison, Amy Kuceyeski, Mert R. Sabuncu

**Affiliations:** School of Electrical & Computer Engineering, Cornell University; Radiology, Weill Cornell Medicine

## Abstract

Task-based and resting-state represent the two most common experimental paradigms of functional neuroimaging. While resting-state offers a flexible and scalable approach for characterizing brain function, task-based techniques provide superior localization. In this paper, we build on recent deep learning methods to create a model that predicts task-based contrast maps from resting-state fMRI scans. Specifically, we propose BrainSurfCNN, a surface-based fully-convolutional neural network model that works with a representation of the brain’s cortical sheet. Our model achieves state of the art predictive accuracy on independent test data from the Human Connectome Project and yields individual-level predicted maps that are on par with the target-repeat reliability of the measured contrast maps. We also demonstrate that BrainSurfCNN can generalize remarkably well to novel domains with limited training data.

## Introduction

Task-based functional magnetic resonance imaging (tfMRI) has been an indispensable tool for probing neural correlates supporting cognitive, emotional and movement-related processes in the human brain. Activation patterns extracted from tfMRI have been used to characterize the functional anatomy of the human brain (***Besle et al., 2013**; **Barch et al., 2013**; **Gordon et al., 2017***), or derive neural biomarkers for individual behavioral measures such as working memory capacity (***McNab and Klingberg, 2008***), visual attention (***Mukai et al., 2007***), loss aversion (***Tom et al., 2007***) or reading ability (***Wang et al., 2019**; **Nijhof and Willems, 2015***). However, tfMRI requires careful design and expensive subject training to elicit the appropriate cognitive components that the experiment intends to investigate (***Church et al., 2010**; **Rosazza et al., 2018***). On the other hand, resting-state fMRI (rsfMRI), which measures spontaneous, slow-changing fluctuations of brain activity in the absence of external stimuli has become the workhorse in a growing number of neuroscience studies, in part due to its ease of acquisition and higher tolerance to confounds (***Power et al., 2014b***; ***Dubois and Adolphs, 2016***). RsfMRI can reveal a wide range of large-scale brain networks and states associated with heterogenous cognitive processes (***Smith et al., 2009***; ***Yeo et al., 2011**; **Power et al., 2011***).Furthermore, resting-state functional connectivity has been demonstrated to yield distinct “fingerprints” unique to individuals (***Finn et al., 2015***; ***Amico and Goñi, 2018***). Despite the differences in methodologies, signals captured by tfMRI and rsfMRI are likely to arise from similar anatomical connections and neural processes, as evidenced by significant overlaps between these two modalities (***Smith et al., 2009***). This suggests that individual task-based brain activity may be predictable from resting-state functional connectivity; indeed, such predictive models based on linear regression have previously been proposed (***Tavor et al., 2016***; ***Cole et al., 2016***). In this work, we revisit this problem using the modern tools of deep learning.

While recent advances in machine learning have enabled dramatic progress in a wide range of fields (***LeCun et al., 2015***), the functional MRI community has been more reluctant to adopt and promote deep *learning* (***Bzdok and Yeo, 2017***). Much of the hesitation in neuroimaging research can be attributed to lack of large-scale high-quality datasets. For example, in computer vision, datasets such as ImageNet (***Russakovsky et al., 2015***) with millions of samples have made training high-capacity neural networks possible. Neuroimaging datasets, today, typically consist of hundreds of subjects or fewer. Furthermore, fMRI data can be highly noisy due to motion or physiological artifacts (***Power et al., 2015***). The relatively low sample size and low SNR regime of neuroimaging makes training high-capacity neural networks exceedingly difficult. Thus, we believe that it is important to implement neural network architectures that take full advantage of the structure of neuroimaging data, while maximizing the available SNR.

In this work, we propose a surface-based neural network called BrainSurfCNN to tackle the problem of predicting subject-specific task contrasts from resting-state functional connectivity. Most neural networks applied to brain imaging operate either in 3D volume, 2D slices, or with region-level vectorized data (e.g. ***Li et al.*** (***2018***); ***Kamnitsas et al.***(***2017***)). In the specific context of functional connectivity, a way of representing similarity of regions’ (ROIs) fMRI time series, common approaches include working with 2D resting-state functional connectivity (rsFC) matrices (***Kawahara et al., 2017***), population-based graphs (***Parisot et al., 2017***), or multi-channel 3D volumes (***Khosla et al., 2019a***), which are treated as inputs for neural networks. Unlike rsFC matrices and population-level graphs, which make use of a low-dimensional representations (pairwise functional connectivity between regions of interest, or ROIs), we use a much richer representation of functional connectivity (vertex-to-ROI). While our method of constructing functional connectivity is closely related to the multi-channel voxel-to-ROI 3D volumes in (***Khosla et al., 2019a***), we work with a surface representation that captures the cortical geometry and allows modeling of fMRI signals on the gray-matter cortical sheet. Furthermore, inter-subject alignment and spatial smoothing on the cortical surface have been shown to better preserve the signal and yield more statistical power for detecting functional activations (***Anticevic et al., 2008***; ***Klein et al., 2010***; ***Frost and Goebel, 2012***). The fMRI field is increasingly recognizing the benefits of surface-based analysis and there has been a substantial shift from volume-based neuroimaging analysis to surface-based ones (***Coalson et al., 2018***), spearheaded by large-scale projects such as the Human Connectome Project (HCP) (***Glasser et al., 2013***). Building on these developments, in this paper we show that the proposed BrainSurfCNN achieves state-of-the-art predictions of individual task contrasts from resting-state functional connectivity, which is on par with the repeat reliability of the contrast signal. Figure 1 shows a representative example of the HCP’s “Social Cognition: Theory of Mind” task contrast for one subject. Thefirsttwo rows are tfMRI derived maps (target and repeat), followed by our BrainSurfCNN model’s prediction of the task contrast based on the resting functional connectivity fingerprint, presented in the third row. Dice scores, commonly used for evaluating accuracy of image segmentation, quantify the overlap with the target contrast map at different thresholds of activation. We observe that our model’s predictions are remarkably consistent with the tfMRI measurements.

**Figure 1.**
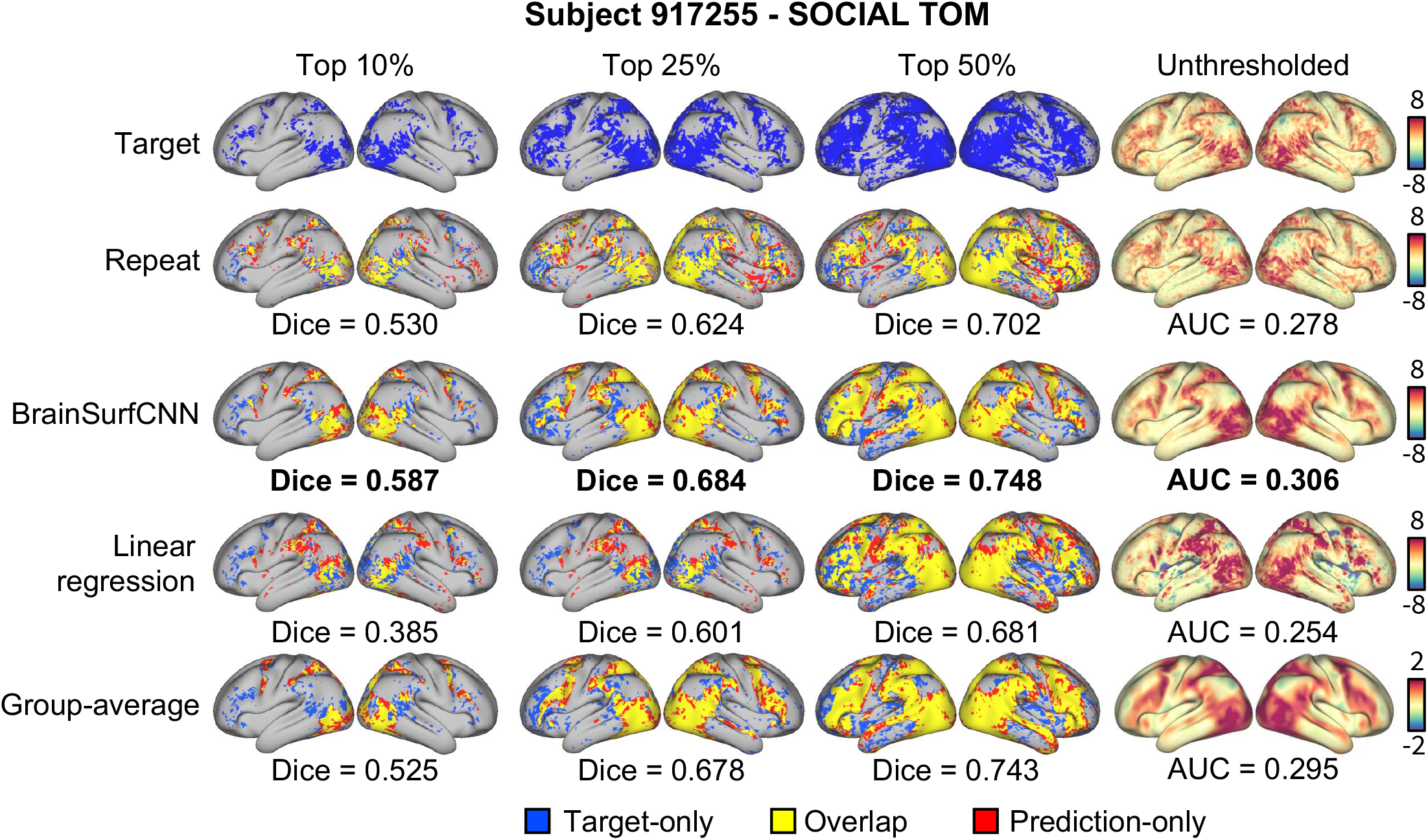
BrainSurfCNN model accurately predicts both coarse and fine-grained features of an individual subject’s task contrast in the HCP dataset. The leftmost three columns show the extent of overlaps between the target thresholded (fMRI-derived) activation map and predicted (from BrainSurfCNN and Linear regression) or reference maps (derived from repeat scan of the same individual performing the same task and group average task contrast). Blue represents the target activation, red represents the prediction or reference, and yellow is the overlap. The rightmost column shows the unthresholded activation maps, Dice overlap is indicated below the corresponding panel. “SOCIAL TOM” is short for “Social Cognition: Theory of Mind”.

In our analyses, we further demonstrate that the BrainSurfCNN model generalizes well to novel tasks and new subjects using a transfer learning paradigm (***Yosinski et al., 2014***); we hypothesize this was possible due to the multi-task learning set-up used for training. Our results show that with the help of transfer and multi-task learning, neural networks applied to neuroimaging can be adapted beyond the original training dataset and generalize well to other contexts in which data might be lacking.

## Results

### Overview of the BrainSurfCNN model

Figure 2 shows the proposed BrainSurfCNN model for predicting task contrasts from resting-state functional connectomes. BrainSurfCNN is based on the U-Net architecture (***Ronneberger et al.,2015***; ***Milletari et al., 2016***) and uses the spherical convolutional kernel (***Chiyu et al., 2019***) to operate on the spherical mesh. The most notable feature in the U-Net architecture is the skip connections that copy features from the encoding arm (outputs from the downstream blocks in Figure 2) to the inputs of the decoding arm (inputs for the upstream blocks in Figure 2). The skip connections were found to improve U-Net predictive accuracy of fine-grained details in U-Net’s original task of image segmentation (***Ronneberger et al., 2015***). Our ablation study (Supplemental Table 1) shows skip connections similarly improves predictive quality in our image generation task. BrainSurfCNN’s input is the functional connectomes, represented as multi-channel data attached to the icosahedral mesh vertices. Each input channel is computed as the Pearson’s correlation between the vertex timeseries and the average timeseries within target ROIs. The ROIs were derived from group-level independent component analysis (ICA) (***Smith et al., 2013***). The input and output surfaces are fs_LR templates (***Van Essen et al., 2012***) with 32,492 vertices (fs_LR 32k surface) per brain hemisphere. The left and right hemispheres are symmetric in the fs_LR atlases, i.e., the same vertex index in both hemispheres corresponds to contralateral analogues. Thus, each subject’s connectomes from the two hemispheres can be concatenated, resulting in a single input icosahedral mesh with the number of channels equalling twice the number of ROIs. BrainSurfCNN’s output is also a multi-channel icosahedral mesh, in which each channel corresponds to one fMRI task contrast. This multi-task prediction setting promotes weight sharing across contrast predictions.

**Figure 2.**
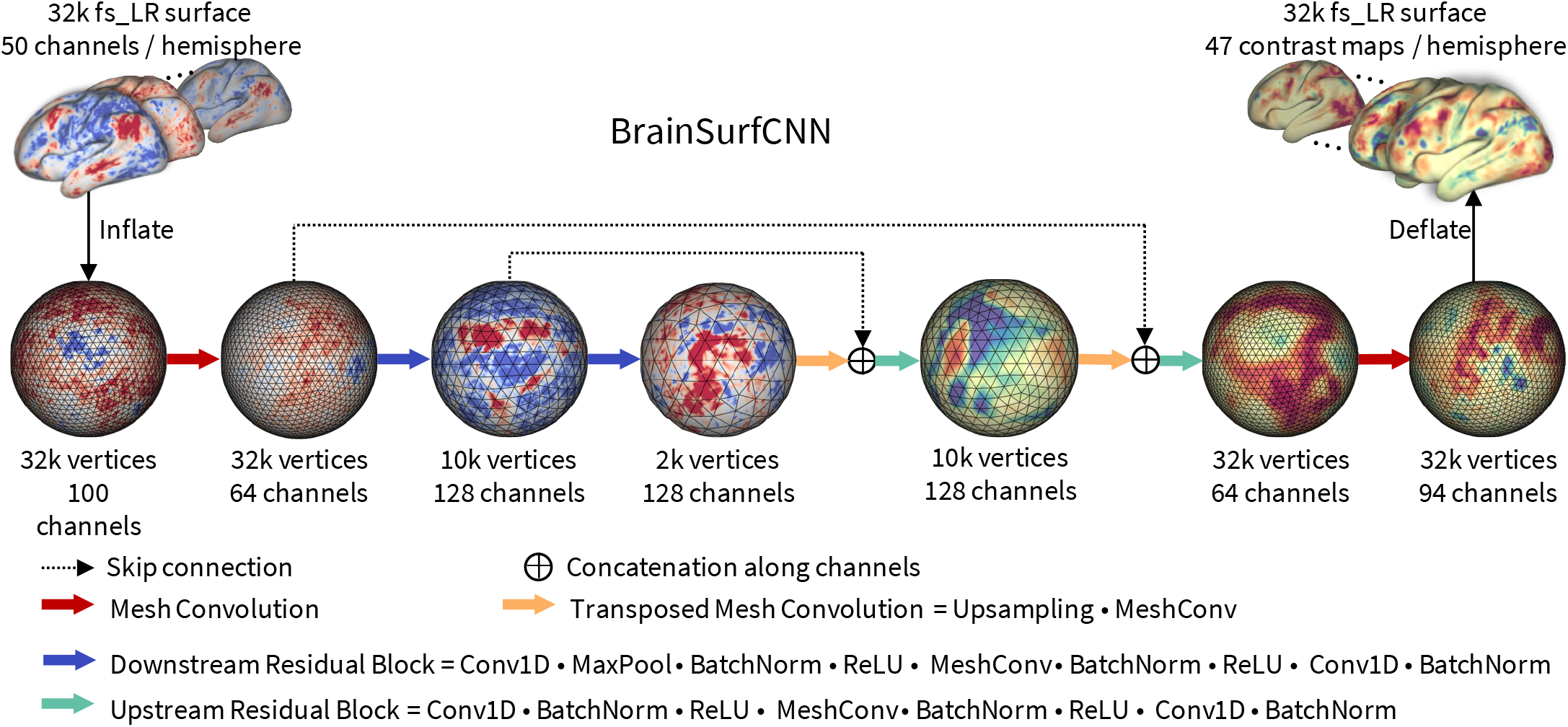
BrainSurfCNN model. BrainSurfCNN is a surface-based fully-convolutional neural network based on the U-Net architecture (***Ronneberger et al., 2015***) with spherical convolutional kernels (***Chiyu et al., 2019***). BrainSurfCNN’s input and output are multi-channel icosahedral fs_LR meshes (***Van Essen et al., 2012***). Each input channel is a functional connectivity feature, measured by Pearson’s correlation between the vertices’ timeseries and the average timeseries of an ROI. Each output channel corresponds to a fMRI task contrast. Details of the model and data formats are found in Section Methods and Materials.

### BrainSurfCNN’s predictive accuracy approaches reliability

We applied BrainSurfCNN to the Human Connectome Project (HCP) dataset to assess the model’s predictive performance. We used Dice scores to evaluate the overlap between the most activated vertices in the target (measured) and predicted task contrasts (see Section Methods and Materials for details); we applied thresholds between 5*%* and 50*%* to identify the most highly activated vertices.

At a lower threshold (e.g., when looking at 5% most activated vertices), the Dice score measures the correspondence of the fine-grained details between the target and predicted contrasts. At higher thresholds (e.g. 50% most activated vertices), this metric quantifies the global agreement of the anatomical distributions. We also computed an approximate integration of Dice scores across all thresholds (between 5 and 50*%*), i.e. the area under the Dice curve (AUC), as a summary measure over all levels of activation and deactivation.

Nine hundred and nineteen subjects were used for training and validation, while the 39 HCP subjects with a repeat scan were used for testing. Two formulations of BrainSurfCNN were evaluated. The multi-contrast BrainSurfCNN was trained to make predictions for all 47 task contrasts simultaneously, while single-task BrainSurfCNN models were trained separately for each task contrast. We compared BrainSurfCNN predictions against two baselines; the group-average task contrasts and a linear regression model (***Tavor et al., 2016***) (details are in Section Methods and Materials). In addition, we used the repeat tfMRI scans to assess the reliability of each subject’s task contrast, which was quantified as the Dice AUC between the maps derived from the target and repeat scans. Out of 47 HCP task contrasts, 24 had a target-repeat reliability AUC higher than the group-average and thus were considered reliably predictable contrasts. Across the 24 reliable task contrasts and test subjects, multi-contrast BrainSurfCNN’s average AUC (0.2834) is on par with the single-contrast BrainSurfCNN models’ average AUC (0.2835), but is higher than the group-average contrasts (0.2693), and the linear model’s baseline predictions (0.2546). BrainSurfCNN also outperforms the average AUC of repeat contrasts (0.2697). As shown in Figure 3, BrainSurfCNN exceeds the target-repeat reliability AUC score for 22 out of 24 reliably predictable task contrasts.

**Figure 3.**
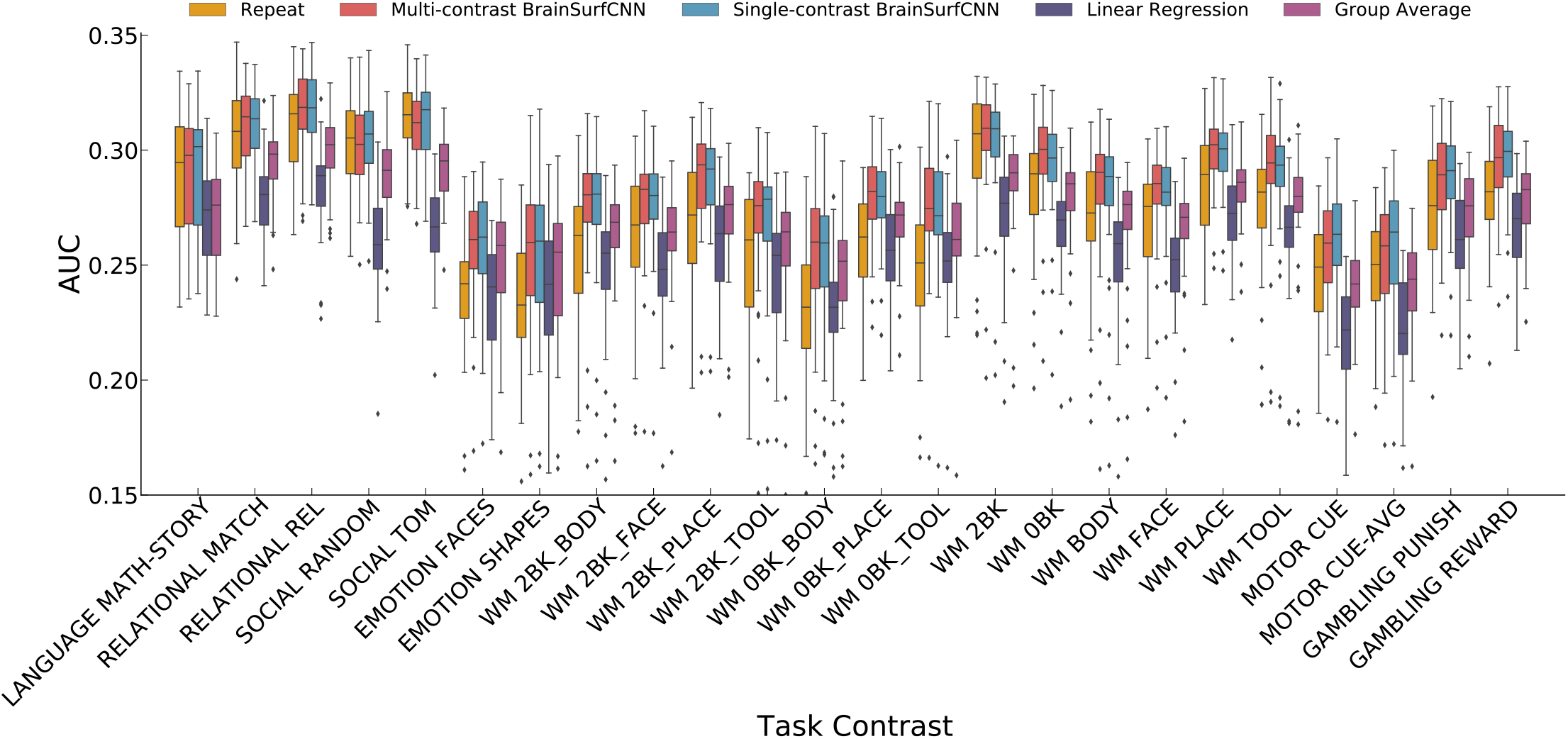
BrainSufCNN prediction is better than the linear regression prediction and group-average contrast map while approaching the noise ceiling (repeat task contrast map) across the most reliably predictable HCP task contrasts (whose target-repeat reliability AUC is higher than the group-average). Quality of prediction is measured as the area under the curve (AUC) of Dice overlap between true and predicted thresholded activation maps (Figure 4). “REL”, “AVG are short for “Relational” and “Average”, respectively. “WM” is short for “Working memory” task; “0BK” and “2BK” are short for “0-back” and “2-back” contrasts of the working memory task, respectively. The list of all 47 HCP task contrasts are availble in Supplemental Table 2. **Figure 3-Figure supplement 1.** AUC scores for all 47 HCP task contrasts.

The three contrasts (from different tasks) with the highest reliability AUC are “SOCIAL TOM” (AUC = 0.312), “RELATIONAL REL” (AUC = 0.310), and “WM 2BK” (AUC = 0.297). Figure 4 shows the Dice score curves for these contrasts. The Dice scores of BrainSurfCNN closely approach the reliability Dice scores across all thresholds, suggesting that the BrainSurfCNN prediction well captures the individual-level variation of task contrasts. Conversely, the agreement between the group-average and measured target task contrasts is lower than the repeat measurement, when only a small fraction of top activated or deactivated vertices are considered, but approaches the reliability score when computed over the majority of vertices. This suggests that the group-average map indeed captures large-scale patterns, but smooths over the individual-level fine details. Supplemental Figure 1 to 3 shows the Dice and AUC for all 47 HCP task contrasts.

**Figure 4.**
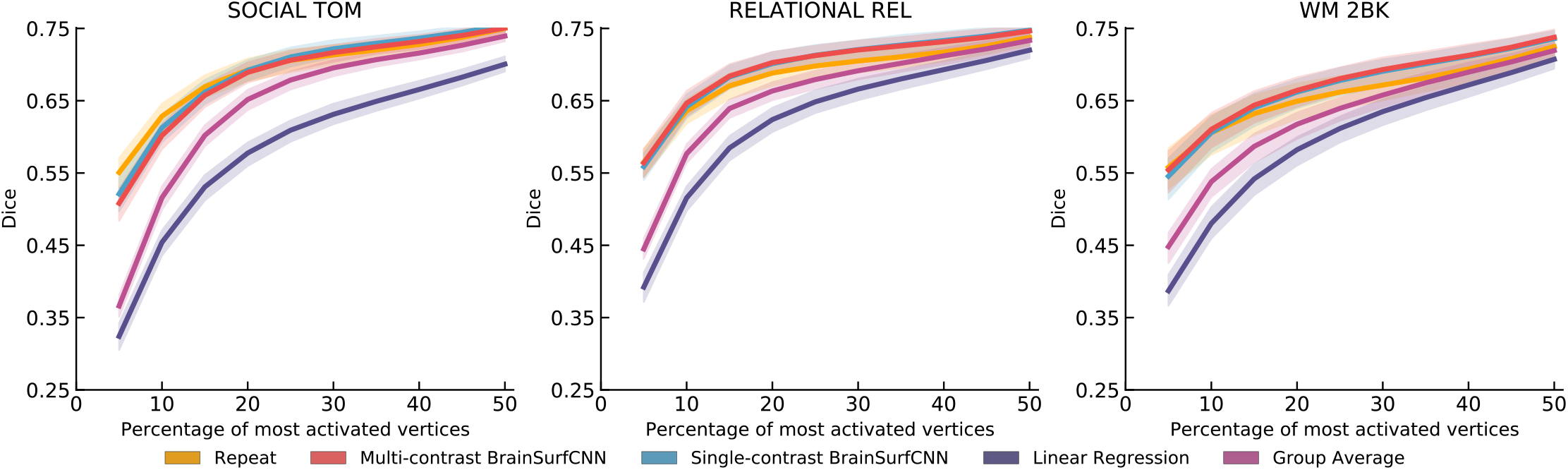
BrainSurfCNN’s predicted task contrasts are comparable to contrasts from repeat task contrasts. For the three most reliable HCP task contrasts (highest overlap with repeat scan), namely “SOCIAL TOM” (social cognition: theory of mind), “RELATIONAL REL” (relational processing), “WM 2BK” (working memory: 2-back), BrainSurfCNN’s Dice overlap with the true activation maps is close to the reliability limit. **Figure 4–Figure supplement 1.** Dice scores for all 47 HCP task contrasts (part 1). **Figure 4–Figure supplement 2.** Dice scores for all 47 HCP task contrasts (part 2). **Figure 4–Figure supplement 3.** Dice scores for all 47 HCP task contrasts (part 3).

### BrainSurfCNN predictions are highly subject-specific

Figure 5 displays the agreement (AUC) across individual subjects’ predicted or repeat measured contrasts (columns) and their measured target contrasts (rows). A subject’s prediction is considered identifiable if it achieves the highest AUC with the subject’s own target contrast, i.e. the diagonal element has the highest value in the column. We compute the identification accuracy as the fraction of subjects that are identifiable. Across the three most reliable task contrasts (SOCIAL TOM, RELATIONAL REL, and WM 2BK), both the repeat scan and BrainSurfCNN predictions yield a 100% identification accuracy, with the linear regression baseline having lower accuracy.

**Figure 5.**
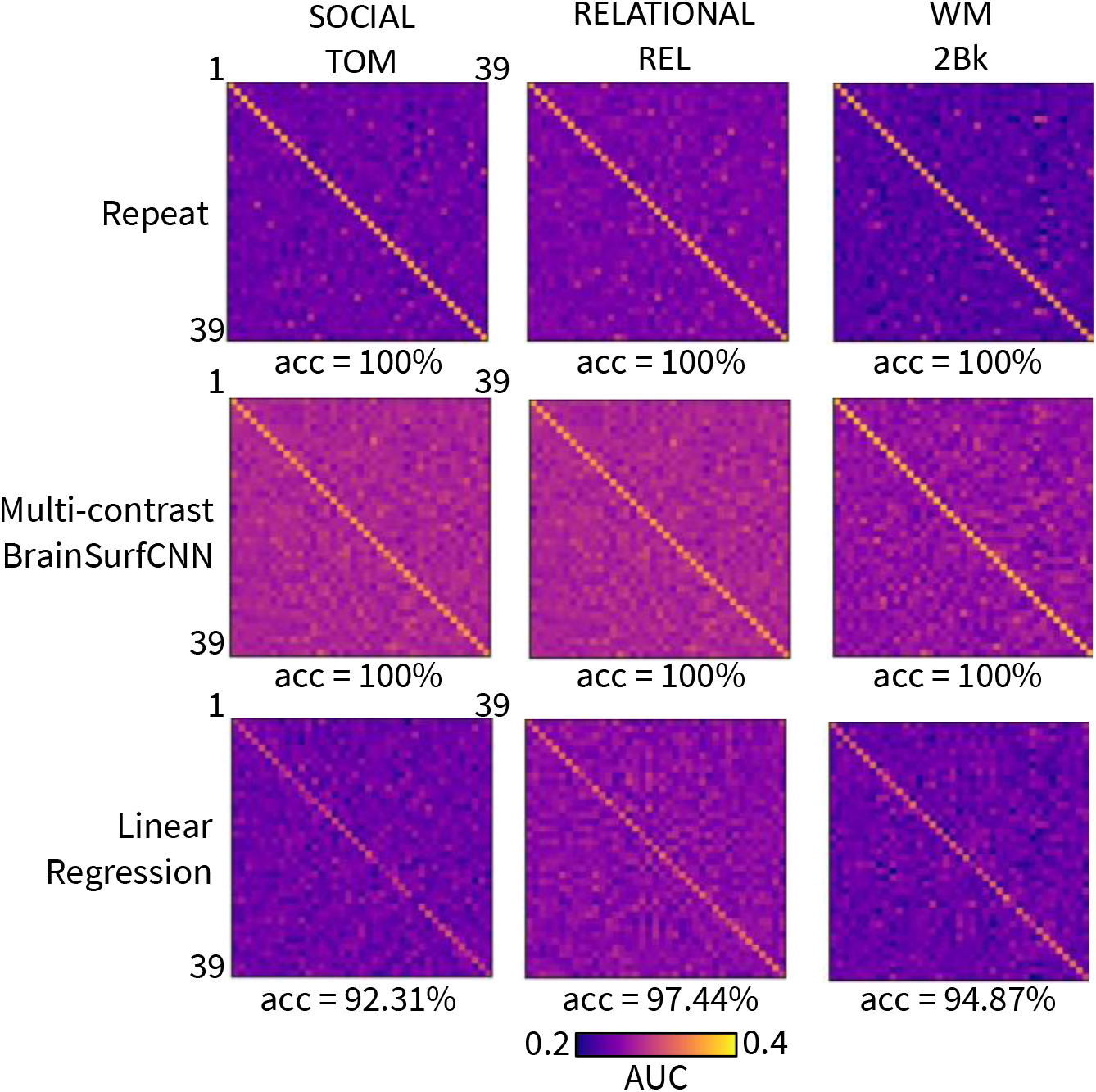
AUC values of prediction versus target measured subject contrasts for 3 most reliable HCP task contrasts, across 39 test subjects. Each row corresponds to a subject’s target task contrast and each column corresponds to the subject’s prediction. The accuracy score below each matrix is the identification accuracy of the model’s prediction for each task.

Figure 6 shows the measured (target) and predicted task contrasts for 3 subjects, including both the unthresholded contrast maps (top rows) and the top 25*%* most activated vertices (bottom rows). The task contrast in consideration is “SOCIAL TOM”, which has the highest reliability. The three subjects, “662551”, “917255”, and “115320” have the 10th, 50th, and 90th percentile “target vs. group-average” AUC among the test subjects, respectively. Thus, these three subjects represent varying degrees of deviation from the typical (group-average) contrast. Focusing on the prefrontal cortex, there are subject-specific activation patterns that appear in the repeat scans. For instance, the replicable activation pattern of the prefrontal cortex in subject “115320” is more laterally dominant, and subject “917255” has more activation in the inferior region of the prefrontal cortex. On the other hand, the activation pattern of subject “662551” is more sparsely distributed in the the prefrontal cortex. Such subject-specific characteristics are captured in the predictions computed by BrainSurfCNN, also reflected by a high Dice overlap between the predicted task contrast and the measured target contrasts for the top 25*%* most activated vertices.

**Figure 6.**
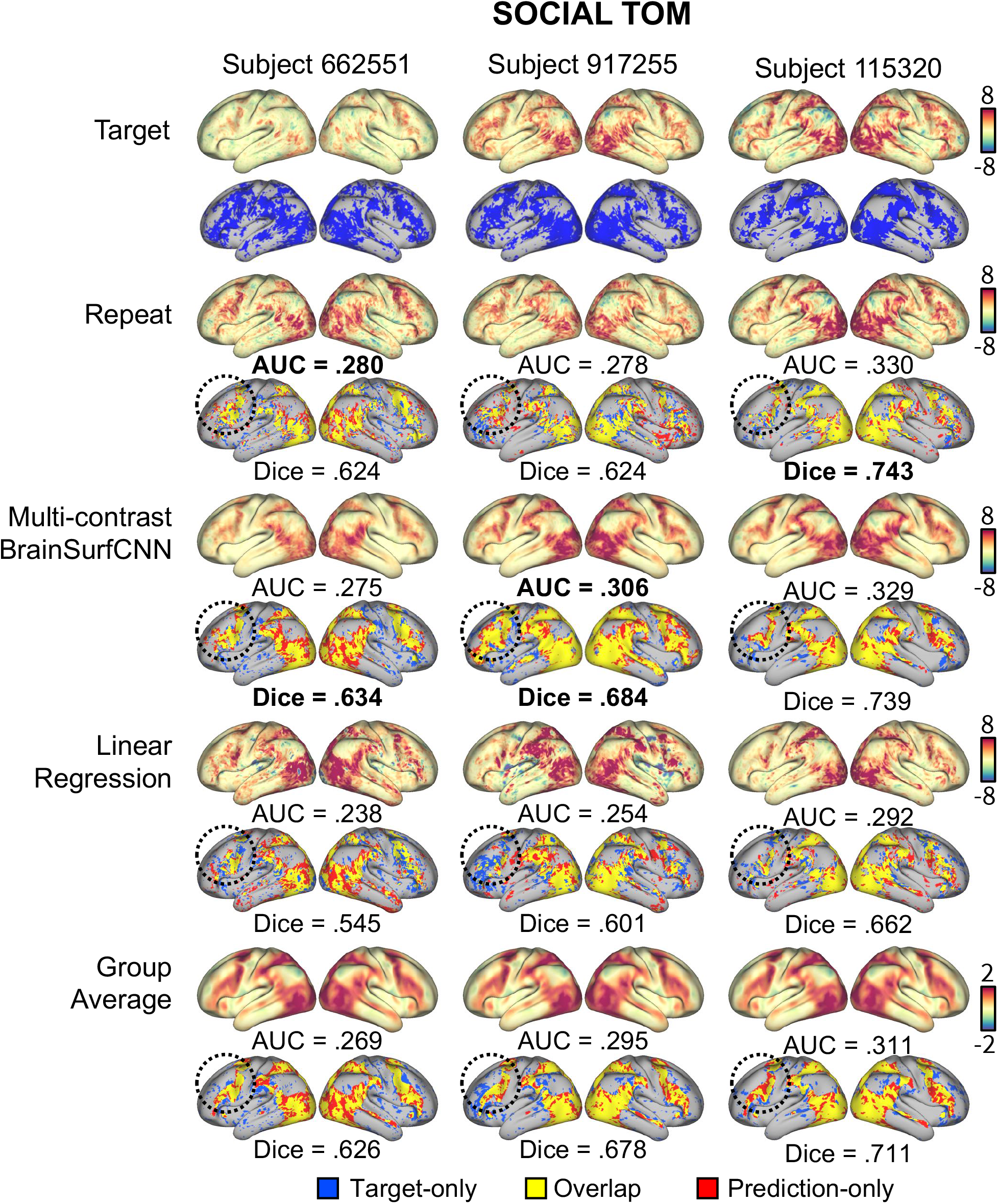
Measured and predicted “SOCIAL TOM” (Social cognition, theory of mind) task contrast for three representative subjects in the HCP dataset. Each row shows both the unthresholded activation maps (top) and thresholded maps of the top 50% most activated vertices (bottom). Blue indicates activation in the measured contrast, red is the predicted or reference activation and yellow is the overlap. The circled areas show activation patterns of the prefrontal cortex distinct to each subject that are replicable in both the repeat contrasts and BrainSurfCNN prediction.

### Transfer learning improves pretrained BrainSurfCNN’s predictive performance on smaller datasets

We investigated BrainSurfCNN’s generalizability beyond the HCP dataset on which the model is originally trained. To do so, we explored model predictions on two Population Imaging of Psychology (PIOP) datasets under the the Amsterdam Open MRI Collection (AOMIC) (***Snoek et al., 2021***)s. Each dataset has both resting-state and task-based fMRI data of healthy participants; PIOP1 has 216 subjects with 5 tasks each and PIOP2 has 226 subjects with 3 tasks each. All AMOIC data, including raw and derivatives, are publicly available (***Snoek et al., 2021***).

There are significant differences between the HCP and AMOIC PIOP datasets, including differences in the task paradigms and scanning procedures. In addition, while the resting-state and task-based fMRI in the HCP dataset was preprocessed directly on the fs_LR surface space (***Van Essen et al., 2013***), the resting-state and task-based fMRI in the PIOP datasets were preprocessed in the volumetric MNI space with fmriprep (***Esteban et al., 2019***). As an extra preprocessing step, resting-sate and task-based fMRI data (t-stats maps) from the PIOP datasets were projected to the fs_LR surface templates via the fsaverage space for all subsequent training and prediction (***Wu et al., 2018***). For both the PIOP1 and PIOP2 datasets, 50 subjects were held out for testing, leaving the rest for training and validation. The partitions were selected to ensure that all training and validation subjects has both resting-state and all task contrasts, but not all test subjects necessarily had all task contrasts.

Two versions of BrainSurfCNN were assessed, one was trained denovo on the training subset of each PIOP dataset using a random intialization - BrainSurfCNN (random init) - while another was finetuned from the HCP-pretrained model using the same PIOP data (BrainSurfCNN finetuned). The group-average task contrasts and linear model predictions were used for comparison.

Figure 7 shows the Dice scores across all activation thresholds and the overall AUC for the 5 task contrasts in the PIOP1 dataset. When training BrainSurfCNN from scratch - BrainSurfCNN (random init) - the model’s prediction was poor; we hypothesize this is due to the relatively small sample size of the PIOP1. On the other hand, prediction quality was greatly improved by finetuning the model pretrained on the HCP dataset, i.e. BrainSurfCNN (finetuned). Figure 8 shows the predicted and measured contrast maps for an average subject and task. The task contrast, “EMOTION MATCHING: EMOTION > CONTROL” has the median target vs. group-average AUC among the 5 PIOP1 task contrasts. For this contrast, subject 0011 has the median AUC with the group-average task contrast. Both Figure 8 and the Dice graphs in Figure 7 show that BrainSurfCNN (finetuned) yields significantly higher Dice scores than the group-average contrasts when the most activated or deactivated vertices are considered. This gap, however, shrinks with more liberal thresholds. Overall, the finetuned BrainSurfCNN model’s predictions are on par or better than the group-average contrasts in terms of AUC, suggesting that the model can capture well the target task contrasts. In this transfer-learning setup, the multi-contrast and single-contrast BrainSurfCNN models adapted well to the new dataset, resulting in similar predictive performance. However, the multi-contrast setup is more computationally efficient (one model for all outputs) relative to maintaining a separate model for each predictive target.

**Figure 7.**
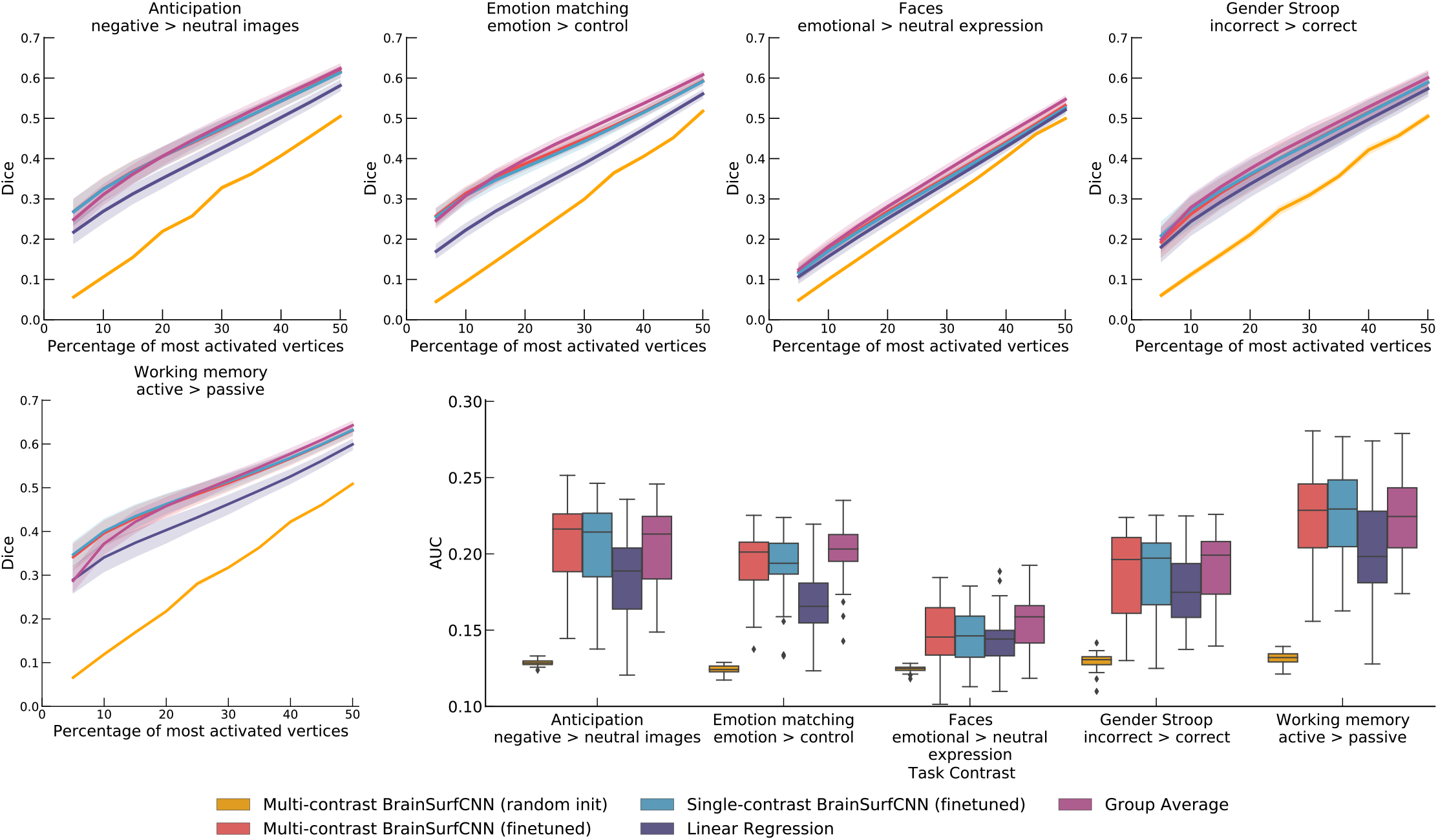
Quality of various model predictions (Dice and AUC) for the Amsterdam PIOP1 dataset. BrainSurfCNN cannot learn effectively if trained from random intialization, resulting in low Dice scores across all thresholds and low overall Dice AUC. However, by finetuning a pretrained model (on HCP data), BrainSurfCNN significantly improves its predictive accuracy and surpasses the linear regression baseline in both the single-contrast and multi-contrast learning setting.

**Figure 8.**
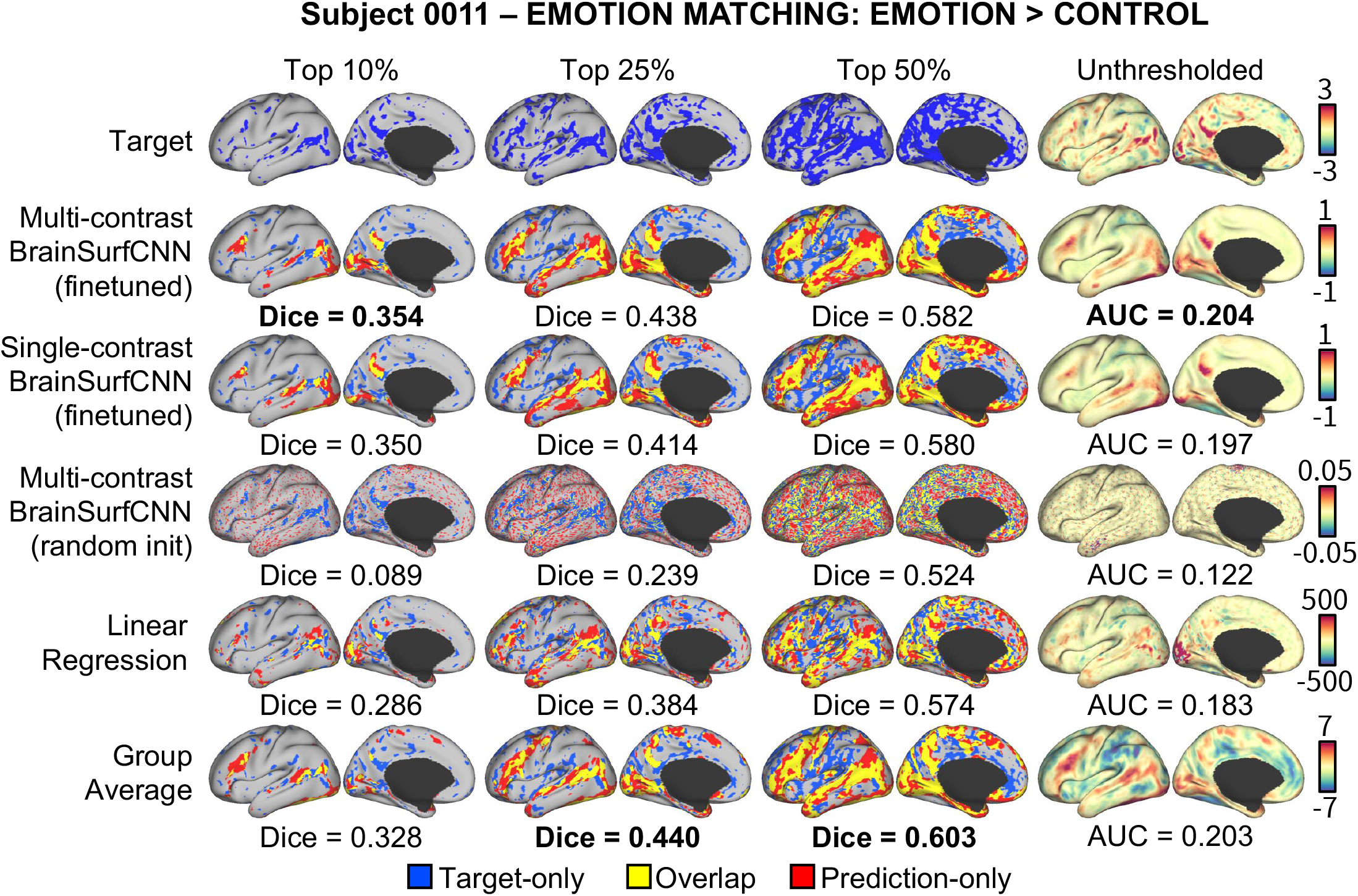
Example set of model predictions for the Emotion Matching: Emotion > Control task for a typical subject in Amsterdam PIOP1 dataset. Without finetuning, BrainSurfCNN trained from random initialization failed to capture meaningful patterns of individual subject’s task contrasts. By finetuning on a model pretrained with HCP data (on a different set of task contrasts), the BrainSurfCNN models were able to predict individual task contrasts in the Amsterdam PIOP1 dataset.

#### Amsterdam OpenfMRI PIOP2

Figure 9 shows the Dice scores across all thresholds and the overall AUC for the 5 task contrasts in the PIOP1 dataset. Similar to our observations above, finetuning the HCP-pretrained BrainSurfCNN model improves predictions over both the linear regression model and the group-average task contrasts. However, in contrast to PIOP1, the BrainSurfCNN trained from random initialization have better predictive performance on PIOP2, possibly because given the same network architecture and roughly the same number of training samples, the model can better learn for a smaller number of predictive outputs (3 task contrasts in PIOP2 compared to 5 task contrasts in PIOP1). Figure 9 shows the predicted and measured contrast maps of the median subject (0014) and the median task contrast (“EMOTION MATCHING: EMOTION > CONTROL”). Similar to the results on the PIOP1 dataset, BrainSurfCNN has the largest gains over the group-contrast for the most activated or deactivated vertices.

**Figure 9.**
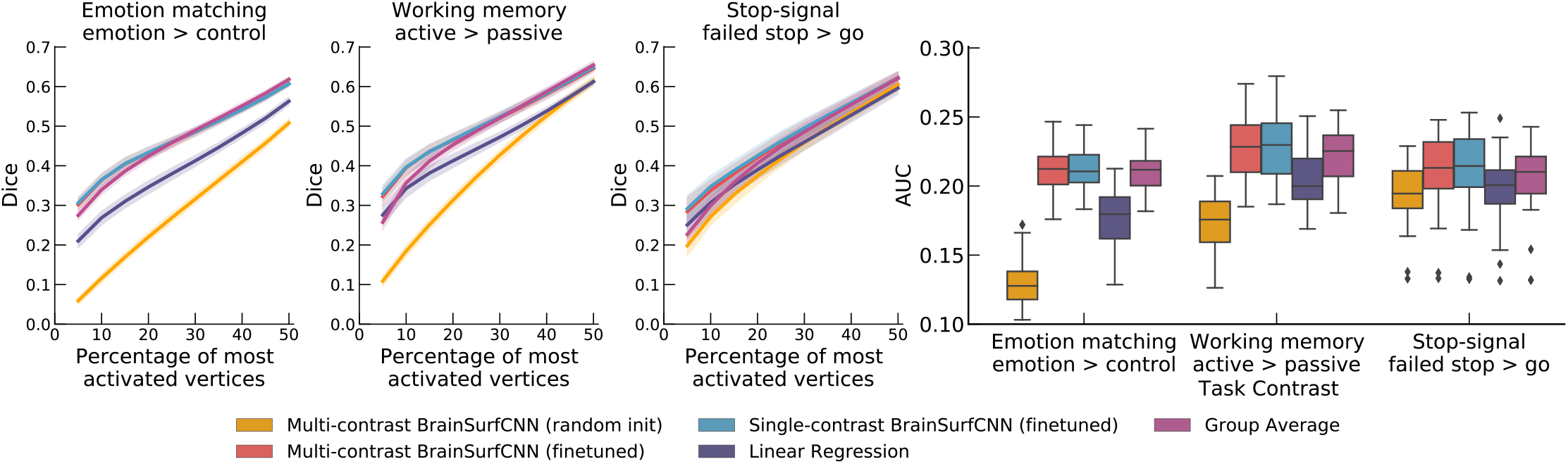
Quality of model prediction (Dice and AUC) for Amsterdam PIOP2 dataset. There is a similar improvement in BrainSurfCNN predictive accuracy via finetuning as for PIOP1 dataset. Compared to the baselines, the finetuned BrainSurfCNN model improves over the linear regression baseline on Dice score across all thresholds, and improves over the group-average reference up to the top 25*%* most activated vertices.

**Figure 10.**
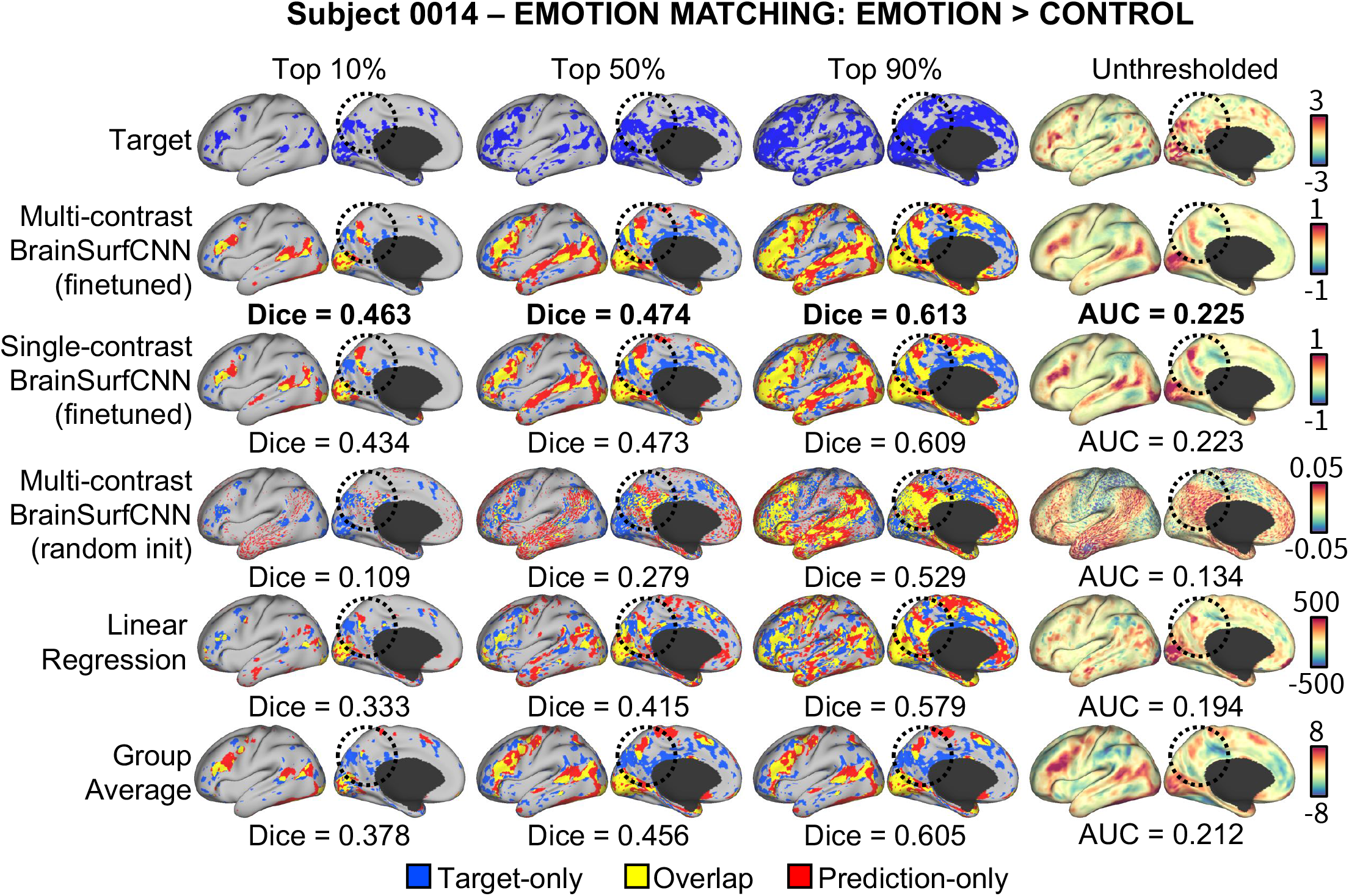
Example set of model predictions for the Emotion Matching: Emotion > Control of a typical subject in Amsterdam PIOP2 dataset. The finetuned BrainSurfCNN could capture both the gross pattern of the task contrast, as well as the subject-specific details. For example, activation in the posterior medial cortex (circled) was predicted correctly by BrainSurfCNN for the given subject that is not significant in the group-average reference.

### The shared representation in multi-task learning enables flexible domain adaptation

We want to investigate the representation shared across predicted task contrasts in the multi-task learning setup of BrainSurfCNN. As all predicted outputs of the multi-contrast BrainSurfCNN model share the same backbone network, which excludes the last deconvolutional layer (Figure 2), we hypothesize that one can achieve domain adaptation via fine-tuning the backbone model using different target contrasts.

We explore this hypothesis by finetuning the backbone of the HCP-pretrained BrainSurfCNN model using a new dataset, the Individual Brain Charting (IBC) (***Pinho et al., 2018**, **2020***). At the time of our analysis (October 2020), the IBC dataset has 12 subjects with both resting-state and task contrast fMRI scans. As the IBC project aims to densely sample cognitive processes, the dataset covers a wide range of task paradigms per subject, including some that are similar but not identical to HCP tasks (see Supplemental Table 2 for task contrasts that are similar between HCP and IBC datasets). For example, IBC “Language” task stimuli are the French translation of the English stimuli used in the original HCP tasks. Together with differences in scanning and preprocessing protocols, there are significant domain shifts between the HCP and IBC datasets even for the same task paradigms. The subjects perform each task in two sessions, one with anterior-posterior (AP) and the other with posterior-anterior (PA) phase encoding during acquisition (***Pinho et al., 2018***). In our experiments, the task contrasts derived from the AP sequence is the measured target, while the contrasts from the PA sequence is treated as the reliability reference (analogous to the repeat contrasts in the HCP dataset). We adapt the BrainSurfCNN model pretrained on the HCP data to the IBC dataset by only finetuning the backbone of the model in a leave-one-task-out procedure. Here the predicted IBC task (which itself consists of multiple contrasts) is treated as unseen data and the remaining HCP contrasts are used for finetuning the BrainSurfCNN backbone. The backbone-finetuned BrainSurfCNN is compared against the multi-contrast and single-contrast BrainSurfCNN, the linear regression models that were only trained on HCP dataset (no finetuning) and the group-average contrasts.

Figure 11 shows the AUC scores across 19 reliable HCP task contrasts in the IBC dataset. Similar to the procedure on the original HCP dataset, reliable contrasts are defined as those whose Dice AUC between the maps derived from the target and repeat scans for HCP task contrasts in the IBC dataset to be higher than the average. The finetuned BrainSurfCNN predictions have better AUC scores than the the repeat contrasts across all but 2 task contrasts (“LANGUAGE STORY” and “SO-CIALTOM”). They also perform better than the pretrained BrainSurfCNN models, linear regression and group-average baselines for all HCP task contrasts. To reemphasize, the finetuned BrainSurfCNN is not trained on the predicted task contrast from the IBC dataset, but merely benefits from the improved backbone finetuned on the other task paradigms. Figure 3 shows that indeed the multi-task setup allows learning shared representation that are beneficial across predictive outputs.

**Figure 11.**
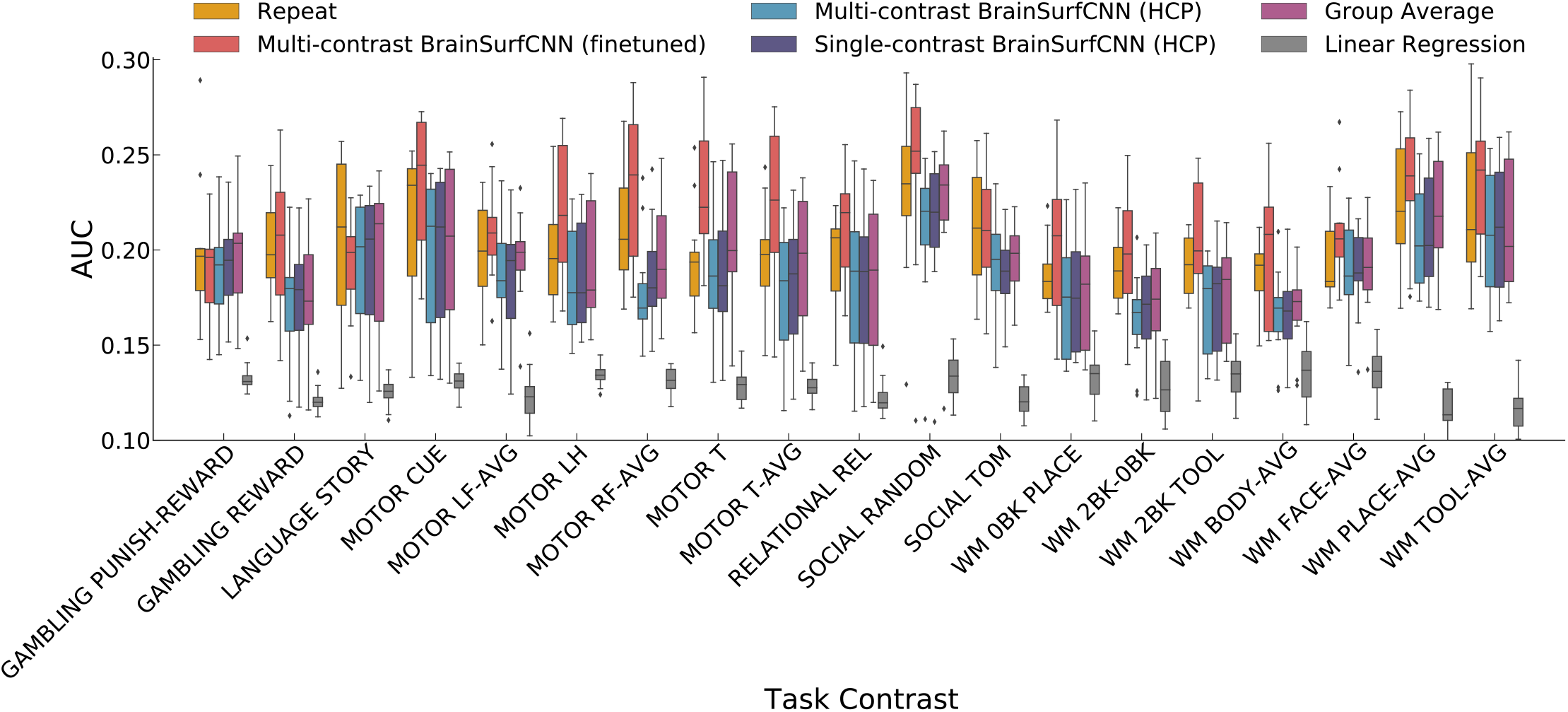
AUC of model predictions on reliable HCP task contrasts in the IBC dataset. By finetuning the multi-contrast BrainSurfCNN’s backbone on other IBC task contrasts, the model improves its predictive accuracy for the test task contrast over the baselines, as well as the non-finetuned models. Note that only the multi-task learning setting allows such leave-one-task-out training procedure without the model’s access to any training samples of the contrast to be predicted in the IBC dataset.

Figure 12 shows the Dice scores of 3 contrasts with the highest average reliability AUC scores from 3 unique tasks. The finetuned BrainSufCNN models improve upon repeat contrasts across all thresholds of activation, except for the top most activated vertices in “RELATIONAL REL” contrast.

**Figure 12.**
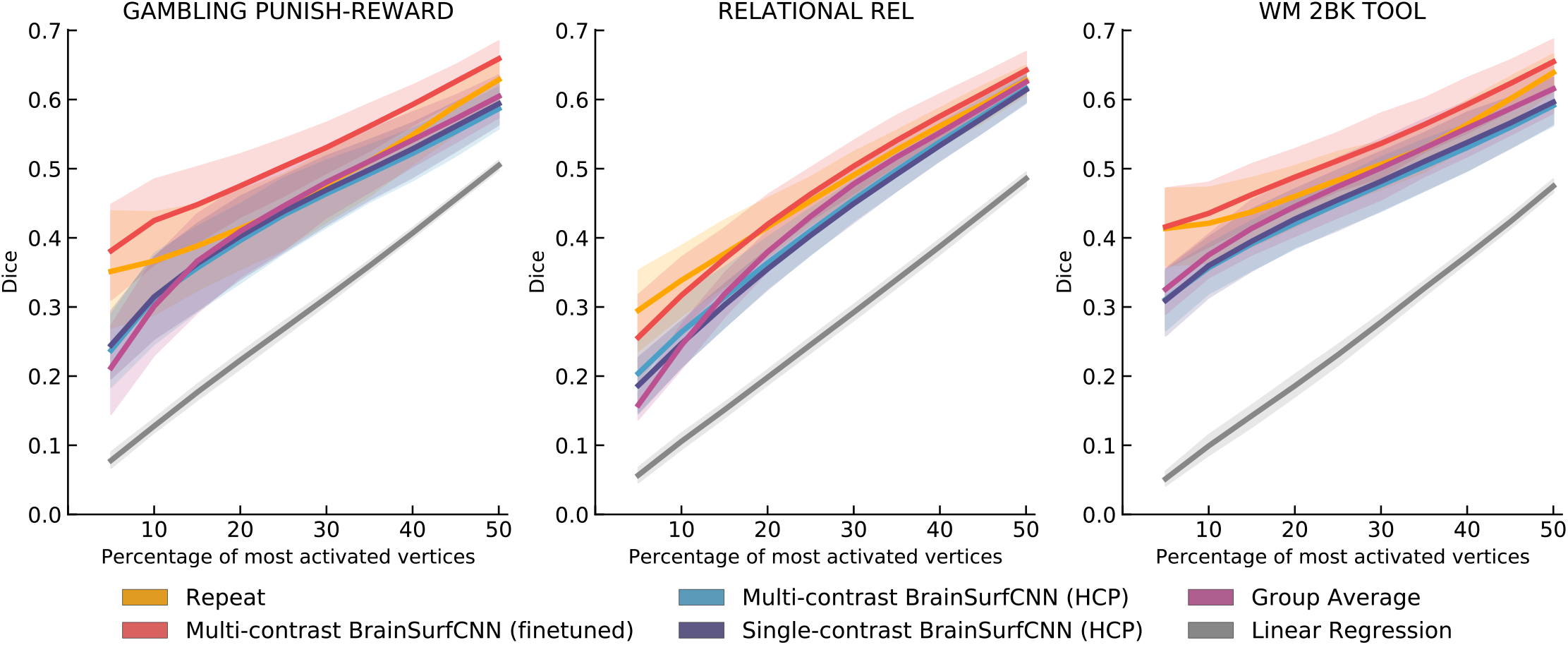
Dice score of model predictions on 3 HCP task contrasts in the IBC dataset at 10th, 50th and 90th average percentile of average AUC among reliable task contrasts. The BrainSurfCNN finetuned on IBC tasks other than the target task greatly improves the pretrained model’s predictive accuracy (in terms of Dice) across all thresholds of activation.

**Figure 13.**
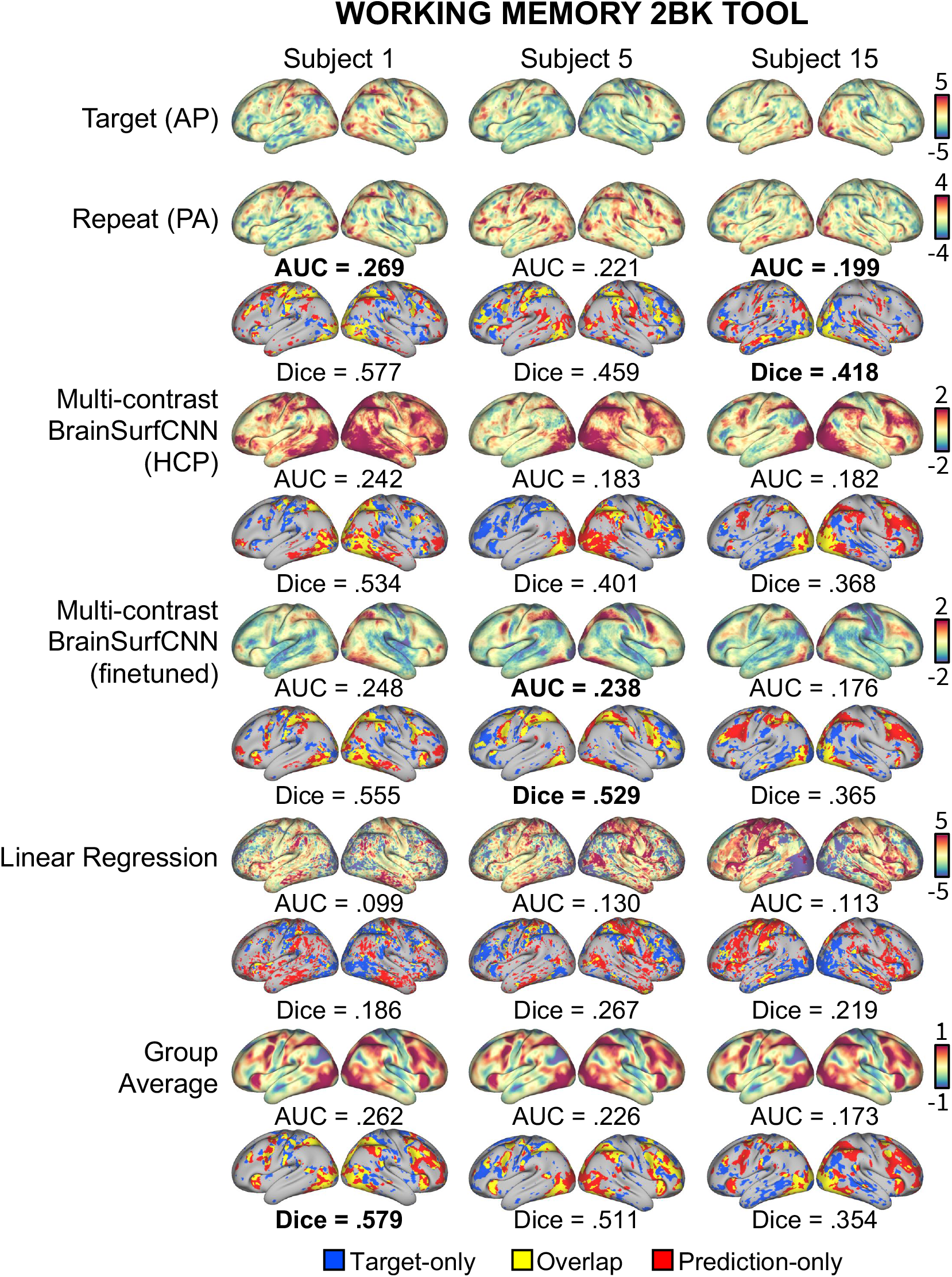
Example model predictions for the “Working memory 2-back tool” contrast of 3 IBC subjects. The finetuned BrainSurfCNN approach the target-repeat reliability in both Dice score for top 50*%* most activated vertices, and overall AUC across all thresholds.

## Discussion

Contrasts derived from task-based fMRI have been instrumental for mapping brain responses across individuals and quantifying how they relate to individual behavioral traits (***McNab and Klingberg, 2008***; ***Mukai et al., 2007***; ***Tom et al., 2007***; ***Wang et al., 2019***; ***Nijhof and Willems, 2015***). Task contrasts are also a useful imaging tool for clinical neurosurgeries (***Matthews et al., 2006***; ***Rosazza et al., 2018***), such as for localizing functional regions, and mapping the impact of lesions. Nonetheless, tfMRI involves meticulous planning and extensive training (***Church et al., 2010***; ***Rosazza et al.,2018***), and can be prohibitive for several subject groups such as some patients or young children. On the other hand, resting-state fMRI is easier to acquire while retaining a rich fingerprint unique to individuals (***Finn et al., 2015***; ***Amico and Goñi, 2018***; ***Tian et al., 2021***) and overlap with task-based fMRI (***Smith et al., 2009***). Therefore, predictive models that can accurately estimate individual-specific task contrasts from the subjects’ resting-state functional connectivity might unlock new venues of research and applications where only rsfMRI is available.

Surface-based preprocessing and analytical methods are increasingly popular in fMRI as they have shown promising results in improving registration, smoothing and functional localization (***Coalson et al., 2018***). In contrast, surfaced-based predictive models for neuroimaging data are relatively less explored. Previous works have used linear regression models to predict vertex-wise brain response for a given task contrast from resting-state functional connectomes (***Tavor et al., 2016***; ***Cole et al., 2016***). In (***Dohmatob et al., 2021***), local gradients estimated from rs-fRMI were used as input features, but the model also made use of a parcel-wise linear regression to predict task activation. Other neural networks-based approaches have used heavily preprocessed inputs, such as low-dimensional ROI-based functional connectivity matrices (***Kawahara et al., 2017***) or population graphs (***Parisot et al., 2017***), which often reduce otherwise rich data into summary metrics and greatly reduce spatial resolution. In a different approach using graph neural networks, (***Zhang et al., 2021***) predicts brain states from fMRI timeseries mapped on to a brain graph. A notable exception that is closely related to our approach is (***Zhao et al., 2019***), which is also a neural network operating on the spherical representation of brain images, but uses a different approximation for the convolutional operation by limiting the kernel to immediate neighbors of each vertex.

In this paper, we present BrainSurfCNN, a neural network model that sets a new benchmark for predicting task activations in individual subjects in the HCP dataset. Furthermore, we demonstrate that using transfer learning, the pretrained BrainSurfCNN model can generalize well to new datasets or new task contrasts that otherwise have limited training samples available. By using a multi-task learning, the model was encouraged to pick out synergies and commonalities shared across tasks that are better learned in unison than isolation (***Caruana, 1997***; ***Baxter, 2000***; ***Maurer et al., 2016***; ***Mensch et al., 2017***), which allows the model to make accurate prediction on task contrasts that were not yet seen during training.

Despite these advances, predicting individual-specific task contrasts is certainly not a resolved challenge. Firstly, inductive bias relevant to characteristics of fMRI data can be introduced to the neural network architecture. For example, if the data is registered to a common brain template, which suggests that the same spherical coordinates correspond to the same anatomical or functional landmarks of the common template, translation-variance bias can be injected to the convolutional operation to improve predictive accuracy, similar to coordinate-aware convolutional kernels’ operation on 2D grid (***Liu et al., 2018***). Secondly, capturing inter-individual variation in model prediction might require new approaches. Training on common loss between a subject’s predicted output and the corresponding measured contrast seems to push the model’s prediction toward the group-average, possibly because inter-subject variation is significantly smaller than the average signal in magnitude, making predicting such small changes challenging. Curiously, learning on individual residuals from the group average, i.e. difference between a subject’s task contrast from the group average, is not effective (not reported). The group average signal seems to act like a regularizer that allows more efficient learning by the neural network. In our experiments with HCP data, by introducing contrastive loss to maximize inter-subject differences of the model outputs, BrainSurfCNN could enhance features specific to each individual subject. Nonetheless, contrastive loss is not effective when finetuning on smaller datasets, possibly because either there are lower signal-to-noise ratios in lower-quality datasets and a more careful search for the right loss margins is needed, or simply because there are an insufficient number of samples for the model to distinguish across samples. Last but not the least, more pre-processing procedures can be experimented with to explore their effect on the downstream predictive task. While we opted for minimally preprocessing all the fMRI data used in our experiments to keep the approach general as a whole, appropriate denoising procedures can change the quality of the input data and improve predictive quality. Furthermore, the limit of 50 ROIs used for computing the input functional connectivity was due to constraints on computing resources. We are working on improving BrainSurfCNN’s efficiency, which can allow using input connectomes with more number of features that can better capture subject-specific variability. To facilitate future studies, the source code for our models and analysis are publicly available at [URL to be available upon publication].

## Methods and Materials

### BrainSurfCNN

BrainSurfCNN’s architecture is based on U-Net (***Ronneberger et al., 2015**; **Milletari et al., 2016***), a fully convolutional neural network originally proposed for segmentation task, and made use of a spherical convolutional kernel (***Chiyu et al., 2019***). The input and output of the model are represented by multi-channel icosahedral meshes of 32,492 vertices (fs_LR 32k surface) (***Van Essen et al.,2012***). Each channel of the multi-channel input mesh is a functional connectivity, measured by the Pearson’s correlation between each vertex’s timeseries and the average timeseries of a target ROI. In our experiments, the target ROIs are parcels derived from group-level 50-component spatial independent component analysis (ICA)(***Smith et al., 2013***). The fs_LR atlases are symmetric between the left and right hemispheres with the same vertex index in both hemispheres correspond to contralateral analogues. Therefore, the two functional connectomes (corresponding to the two hemispheres) of each subject can be concatenated, resulting in a single input icosahedral mesh with twice the the number of channels as the number of ROIs. For BrainSurfCNN’s output, each channel of the multi-channel icosahedral output mesh corresponds to one fMRI task contrast map. By predicting multiple task contrasts simultaneously, weight sharing is promoted across model outputs.

### Reconstructive-contrastive loss

In training BrainSurfCNN, we minimize a reconstructive-contrastive loss, which we describe here. Given a mini batch of *N* samples *B* = {**x**_*i*_}, in which **x**_*i*_ is the target multi-channel contrast image of subject *i*, let 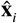 denote the corresponding prediction. The reconstructive-contrastive loss (R-C loss) is given by:

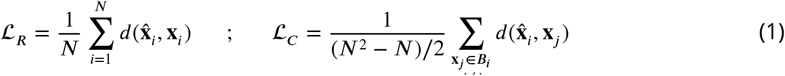

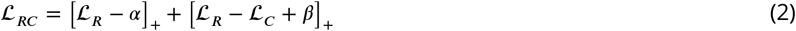

where *d*(.) is a loss function (e.g. *l*^2^-norm). 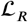, *α* are the same-subject (reconstructive) loss and margin, respectively. 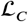, *β* are the across-subject (contrastive) loss and margin, respectively. The combined objective 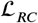 encourages the same-subject error 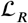 to be within *α* margin, while pushing the across-subject difference 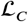 to be large such that 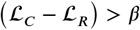.

### Data

#### Human Connectome Project (HCP)

For our benchmarking experiments with a large dataset, we used the minimally pre-processed, FIX-cleaned 3-Tesla resting-state fMRI (rsfMRI) and task fMRI (tfMRI) of 1,200 subjects from the Human Connectome Project (HCP). The dataset’s acquisition and preprocessing were described in (***Glasser et al., 2013**; **Smith et al., 2013**; **Barch et al., 2013***). rsfMRI data was acquired in four 15-minute runs, each with 1,200 time-points per subject. Group-level parcellations derived from spatial ICA were also released by HCP. We used ROIs from the 50-component parcellation for computing the functional connectomes. HCP’s tfMRI data comprises of 86 contrasts from 7 task domains (***Barch et al.,2013***), namely: WM (working memory), GAMBLING, MOTOR, LANGUAGE, SOCIAL RELATIONAL, and EMOTION. Following (***Tavor et al., 2016***), redundant negative contrasts were excluded, resulting in 47 unique contrasts. Out of 1,200 HCP subjects, 46 subjects also have repeat (second visit) 3T fMRI data. Including only subjects with all 4 rsfMRI runs and 47 tfMRI contrasts, our dataset comprised of 919 subjects for training/validation, and 39 independent subjects (with repeat scans) for evaluation.

#### Amsterdam Open MRI Collection (AOMIC)

AOMIC is a collection of multimodal brain imaging datasets from a large number of subjects (***Snoek et al., 2021***). For our experiments with transfer learning, we used two AOMIC datasets: PIOP1 and PIOP2 (PIOP stands for Population Imaging of Psychology). PIOP1 consists of 6 minutes of rsfMRI (480 timepoints at 0.75-second TR) and tfMRI measured from five tasks, namely “Emotion matching”, “Gender Stroop”, “Working memory”, “Face perception”, and “Anticipation”, for 216 subjects. PIOP2 consists of 8 minutes of rsFMRI (240 timepoints at 2-second TR)and tfMRI collected from three tasks, namely “Emotin matching”, “Working memory” and “Stop signal”, for 226 subjects. AOMIC data are organized according to the Brain Imaging Data Structure (BIDS) (***Gorgolewski et al.,2016***).

#### Individual Brain Charting (IBC)

To demonstrate the flexible domain adaptation multi-task learning affords, we used data from the Individual Brain Charting (IBC) project ***Pinho et al*.**(***2020***). IBC dataset includes fMRI data from 12 subjects and 180 task contrasts, 43 of which are also studied in the HCP. IBC data is also organized according to BIDS (***Gorgolewski et al., 2016***).

#### fMRI preprocessing and volume-to-surface projection

Resting-state fMRI data from PIOP and IBC datasets were preprocessed using the FMRIPREP version stable (***Esteban et al., 2019***), a Nipype (***Gorgolewski et al., 2011***) based tool. Each T1w (T1-weighted) volume was corrected for INU (intensity non-uniformity) using N4BiasFieldCorrection v2.1.0 (***Tustison et al., 2010***) and skull-stripped using antsBrainExtraction.sh v2.1.0 (using the OASIS template). Brain surfaces were reconstructed using recon-all from FreeSurfer v6.0.1 (***Dale et al., 1999***), and the brain mask estimated previously was refined with a custom variation of the method to reconcile ANTs-derived and FreeSurfer-derived segmentations of the cortical gray-matter of Mindboggle (***Klein et al., 2017***). Spatial normalization to the ICBM 152 Nonlinear Asymmetrical template version 2009c (***Fonov et al., 2009***) was performed through nonlinear registration with the antsRegistration tool of ANTs v2.1.0 (***Avants et al., 2008***), using brain-extracted versions of both T1w volume and template. Brain tissue segmentation of cerebrospinal fluid (CSF), white-matter (WM) and gray-matter (GM) was performed on the brain-extracted T1w using fast (***Zhang et al., 2001***) (FSL v5.0.9).

Functional data was slice time corrected using 3dTshift from AFNI v16.2.07 (***Cox, 1996***) and motion corrected using mcflirt (FSL v5.0.9 (***Jenkinson et al., 2002***)). This was followed by co-registration to the corresponding T1w using boundary-based registration (***Greve and Fischl, 2009***) with six degrees of freedom, using bbregister (FreeSurfer v6.0.1). Motion correcting transformations, BOLD-to-T1w transformation and T1w-to-template (MNI) warp were concatenated and applied in a single step using antsApplyTransforms (ANTs v2.1.0) using Lanczos interpolation.

Physiological noise regressors were extracted applying CompCor (***Behzadi et al., 2007***). Principal components were estimated forthe two CompCor variants: temporal (tCompCor) and anatomical (aCompCor). A mask to exclude signal with cortical origin was obtained by eroding the brain mask, ensuring it only contained subcortical structures. Six tCompCor components were then calculated including only the top 5% variable voxels within that subcortical mask. For aCompCor, six components were calculated within the intersection of the subcortical mask and the union of CSF and WM masks calculated in T1w space, after their projection to the native space of each functional run. Frame-wise displacement (***Power et al., 2011***) was calculated for each functional run using the implementation of Nipype. Many internal operations of FMRIPREP use Nilearn (***Abraham et al., 2014***), principally within the BOLD-processing workflow. For more details of the pipeline see https://fmriprep.readthedocs.io/en/stable/workflows.html. Preprocessed volumetric PIOP and IBC daa were projected to the fs_LR surface template via fsaverage (***Wu et al., 2018***).

### Baseline

#### Linear regression

The linear regression baseline was implemented according to (***Tavor et al., 2016***), as given by:

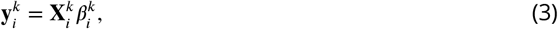

where 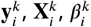 are the vectorized activation pattern, input features, and regressor of the *k*-th parcel in the *i*-th subject, respectively. The 50-component parcellation derived from ICA, provided by HCP was used to compute the linear regression model. 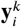 is a vector of length *n_k_* - the number of vertices in the *k*’th parcel in both hemispheres. 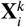 is a *n_k_* × *M* functional connectivity matrix, where each element was computed as the Pearson’s correlation between a vertex and the average timeseries of each of the *M* ROIs (same timeseries used to compute BrainSurfCNN’s input). Following (***Tavor et al., 2016***), a linear model was fit for every parcel and every task of each training sample. These fitted linear models were averaged across all training samples to yield a single predictive model per parcel.

#### Group-average contrasts

Different degrees of inter-subject variability manifest in different task contrasts. Such variability in prediction was a subject of interest for our study. Therefore, we used the group averages as a naive baseline. The group-average task contrasts’ correspondence with individual contrasts would be low/high for tasks with high/low inter-subject variance.

#### Repeat contrasts

We used repeat tfMRI scans (when available) to quantify the reliability of the target contrast maps and evaluate the predictive performance of BrainSurfCNN and the baselines. The repeat contrasts were compared to the first contrasts both in terms of overall correspondence (measured with Dice) and in the subject identification task. We consider these reliability results as an effective upper-bound on performance.

### Experimental setup

#### Data Augmentation and Test-time Ensembling

In the HCP dataset, each subject has 4 rsfMRI runs with 1200 time-points each. As stable functional connectomes can be estimated from fewer than 1200 time-points (***Finn et al., 2015***), we computed a functional connectome from each contiguous half (600 time-points) of every run, resulting in 8 input samples per subject. During BrainSurfCNN training, one connectome was randomly sampled for each subject. Thus, the model was presented with 8 slightly different samples per subject. At test time, 8 predictions were made for each subject and then averaged for a final prediction. For the AOMIC dataset, 4 functional connectomes were computed for each subject from 4 contiguous segments of 120 timepoints. For the IBC dataset, 2 connectomes were computed for each subject, each connectome was estimated from a randomly sampled contiguous segments of 600 timepoints.

#### Training schedule

On the HCP dataset, BrainSurfCNN was first trained for 50 epochs with a batch size of 2 with mean squared error (MSE), i.e. reconstructive loss *L_R_* in Eq.2, using Adam optimizer. Upon convergence, the average reconstructive loss *L_R_* and *L_C_* were estimated from all training subjects, and used as initial values for the margins *α* and *β* in Eq.2 respectively. The initialization procedure forces the model to not deviate from the existing reconstructive error while optimizing for the contrastive loss. Training then continued for another 50 epochs, with the same-subject margin *α* halved and across-subject margin *β* doubled every 20 epochs, which encourages the model to refine further overtime. For transfer learning experiments with the AOMIC and IBC datasets, the best checkpoint of the model when training on HCP dataset with MSE was used as the initialization for finetuning. The finetuning was conducted with MSE as the objective function as the L-R loss did not seem to be sufficiently robust on smaller datasets.

#### Evaluation Metrics

Dice score (***Dice, 1945***) is used to measure the extent of overlap between a predicted contrast and the target contrast map for a given percentage of most activated vertices. At a given threshold of *x%*, Dice score is computed as:

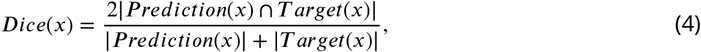

where |*Prediction*(*x*)| denotes the number of top *x%* most activated vertices in the predicted contrast map, |*Target*(*x*)| denotes the number of top *x%* most activated vertices in the contrast, and |*Prediction*(*x*)∩*Target*(*x*)| denotes the number of vertices that overlap between the predicted and target map at the given threshold. By integrating Dice scores over a range of thresholds (e.g. 5*%* to 50*%* most activated vertices), we produce a summary measure - area under the Dice curve (Dice AUC) - for the quality of a model prediction.

#### Quantifying Identification Accuracy

Dice AUCs were computed between the models’ predicted individual task contrast maps and the tfMRI-derived target contrast maps of all subjects. This results in a 39 by 39 AUC matrix for each contrast, where each entry is the Dice AUC between a subject’s predicted contrast (column) and a target contrast map (row), of same or another subject. The diagonal values (Dice AUC between the predicted and target contrast map of the same subject) therefore quantify the (within subject) predictive accuracy for a given contrast. The difference between diagonal and average off-diagonal values (Dice AUC between a subject’s predicted contrast map with the target contrast map derived from another subject’s tfMRI) indicates how much better one subject’s prediction corresponds with the subject’s own tfMRI-derived contrast compared to other subject contrasts. In other words, the *i*-th subject is identifiable among all test subjects using the predicted contrast if the *i*-th element of the *i*-th row has the highest value. For a given task contrast and prediction model, we compute subject identification accuracy as the fraction of subjects with a maximum at the diagonal.

## Acknowledgements

This work was supported by NIH grants R01LM012719 (MS), R01AG053949 (MS), R21NS10463401 (AK), R01NS10264601A1 (AK), the NSF NeuroNex grant 1707312 (MS), the NSF CAREER 1748377 grant (MS) and Jacobs Scholar Fellowship (GN).

---

**Figure 3-Figure supplement 1.**
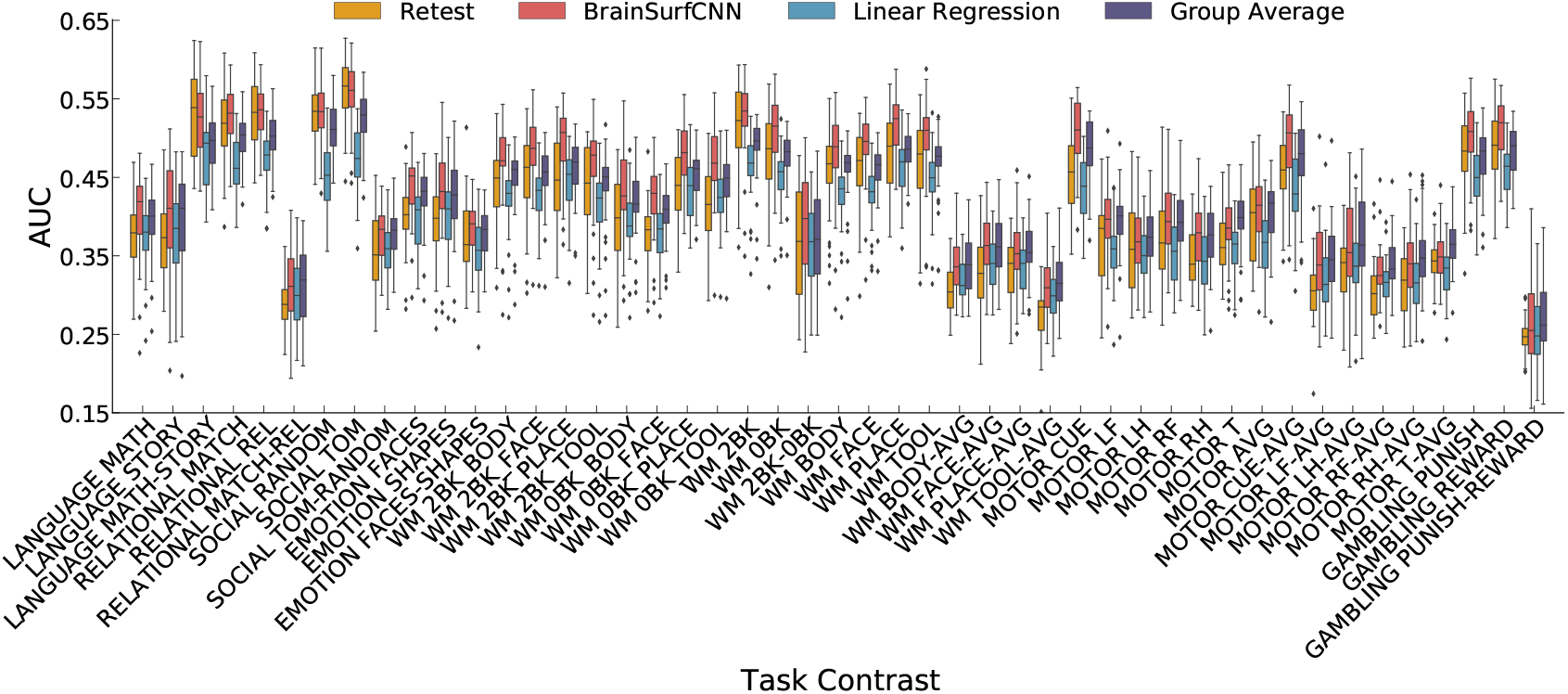
AUC scores for all 47 HCP task contrasts.

**Figure 4–Figure supplement 1.**
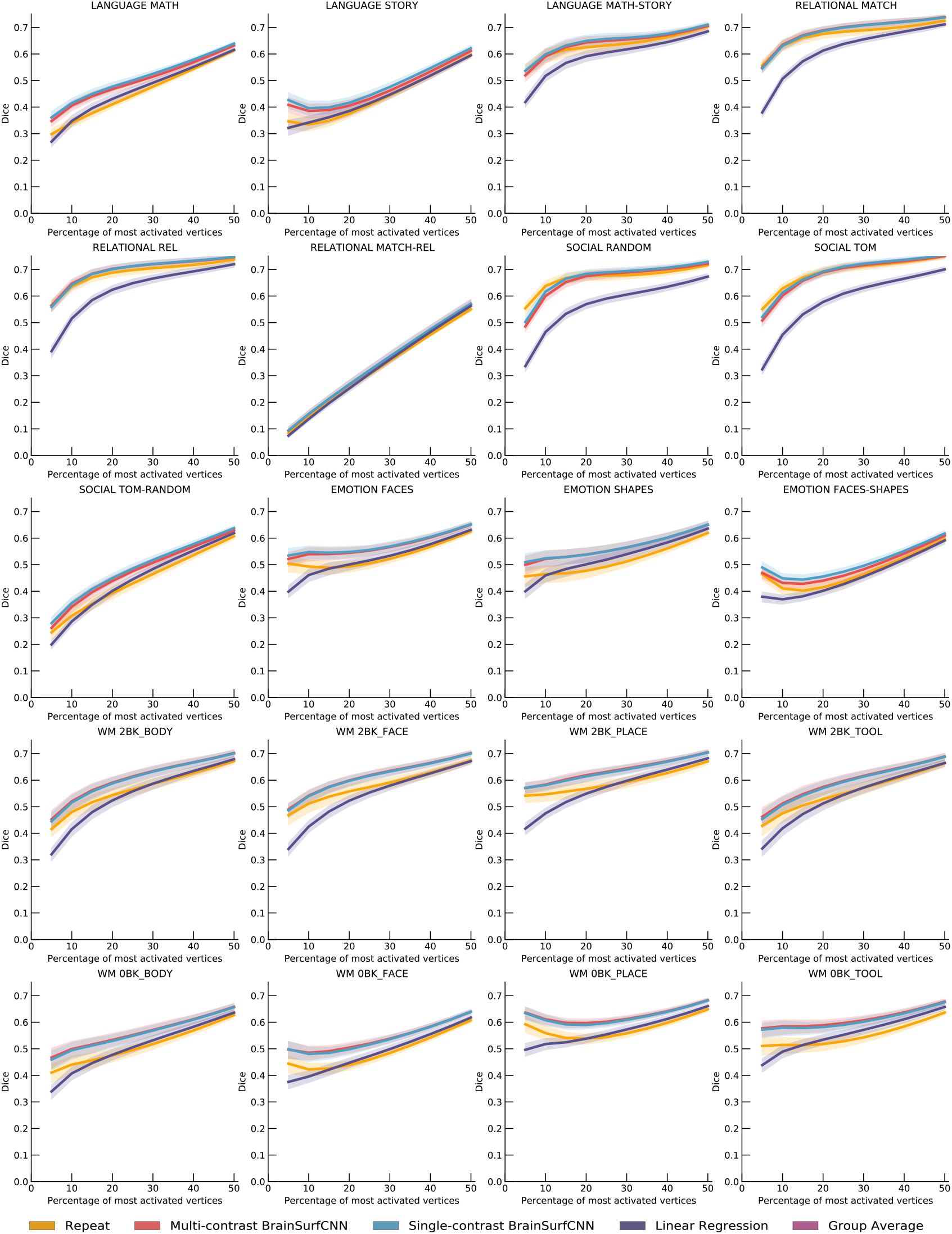
Dice scores for all 47 HCP task contrasts (part 1).

**Figure 4-Figure supplement 2.**
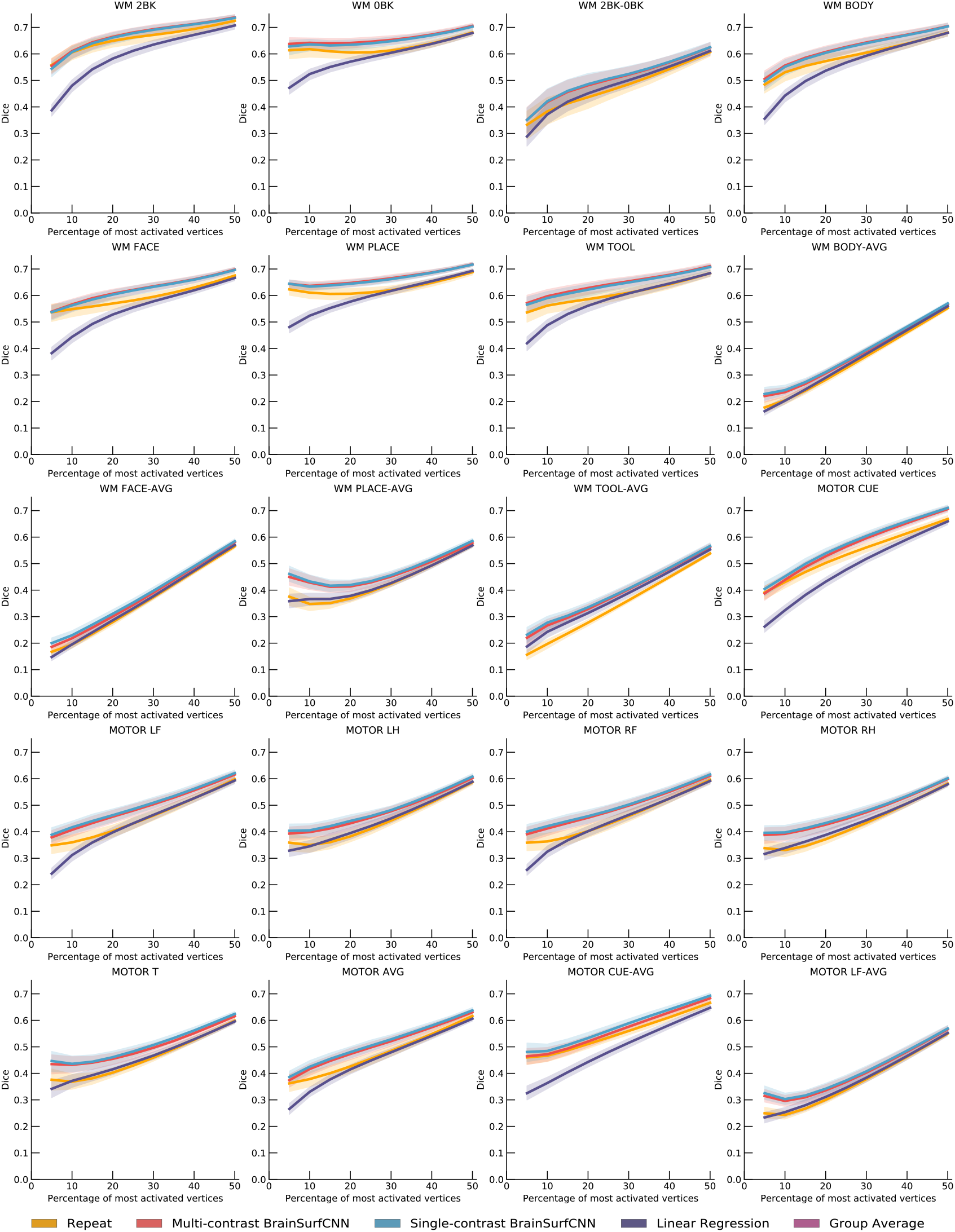
Dice scores for all 47 HCP task contrasts (part 2).

**Figure 4–Figure supplement 3.**
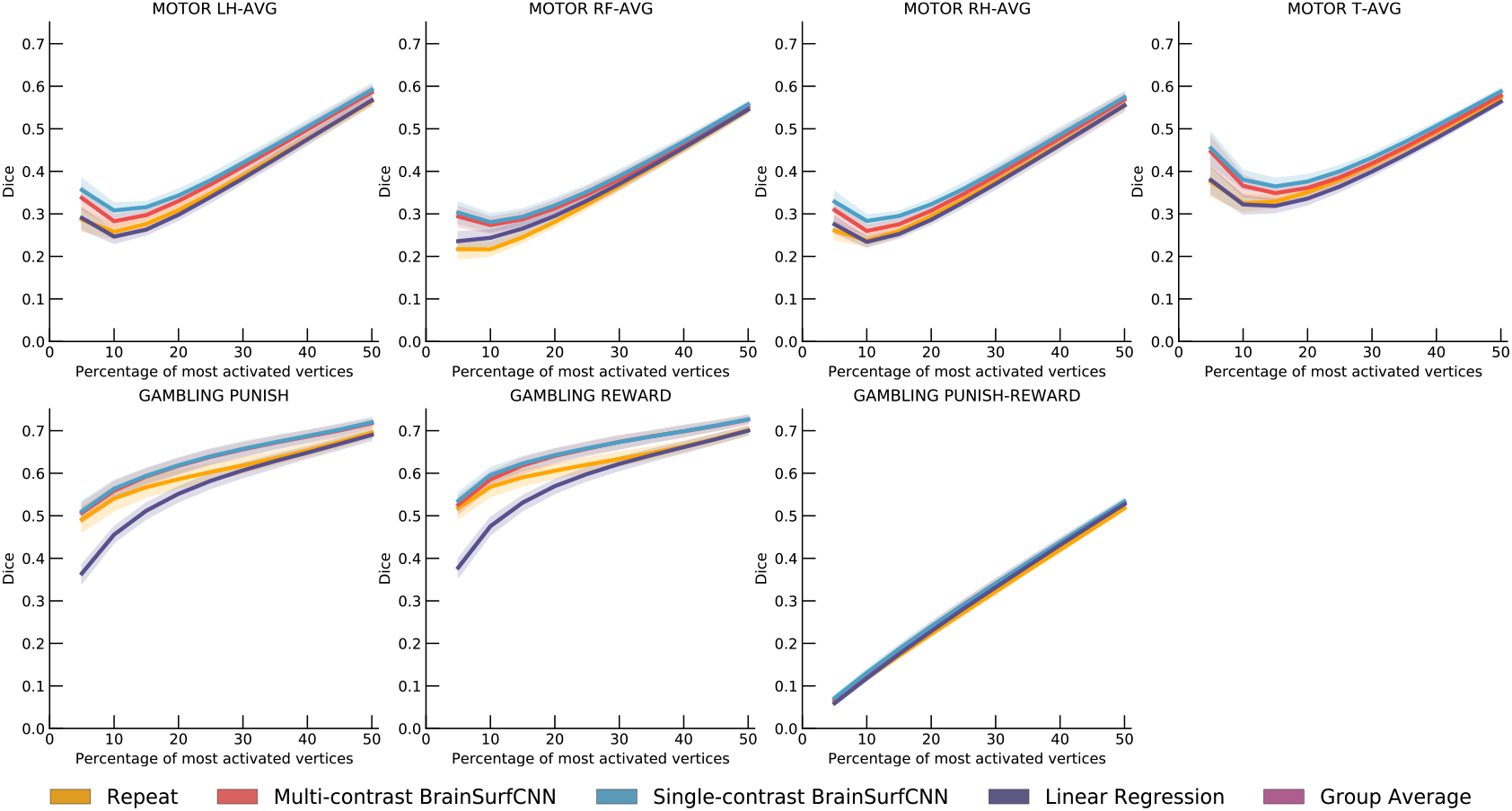
Dice scores for all 47 HCP task contrasts (part 3).

