## Supplemental file for "Predicting Individual Task Contrasts From Resting-state Functional Connectivity using a Surface-based Convolutional Network"

| Model | AUC between prediction and HCP target contrasts |
|---|---|
| BrainSurfCNN | 0.242 ± 0.054 |
| BrainSurfCNN without contrastive loss | 0.241 ± 0.054 |
| BrainSurfCNN without nonlinearity | 0.223 ± 0.052 |
| BrainSurfCNN without skip connections | 0.212 ± 0.054 |

**Supplemental Table 1: ablation study of BrainSurfCNN architecture and training.** The four models used for comparison are (1) full BrainSurfCNN model and training model as described in "Methods and Materials" section; (2) BrainSurfCNN trained only on mean squared error (BrainSurfCNN without contrastive loss) (3) BrainSurfCNN with no nonlinear operation (BrainSurfCNN without nonlinearity) (4) BrainSurfCNN with no skip connection, which is equivalent to an auto encoder architecture, but doubled in number of channels per intermediate layers to retain comparable number of parameters to the full model. The models were evaluated on the HCP test set across all 47 task contrasts using Dice AUC metrics using the checkpoint with the lowest error on the validation set. The ablation study shows that all components of the BrainSurfCNN contributes to the quality of the model prediction.¶

| **Task** | **Contrast** | **Abbreviation** | **Present in IBC dataset** |
|---|---|---|---|
| Language | Math | LANGUAGE MATH | Yes |
| | Story | LANGUAGE STORY | Yes |
| | Math - story | LANGUAGE MATH-STORY | Yes |
| Relational Processing | Relation | RELATIONAL REL | Yes |
| | Match | RELATIONAL MATCH | Yes |
| | Match - relation | RELATIONAL MATCH-REL | |
| Social Cognition | Mental interaction | SOCIAL TOM | Yes |
| | Random interaction | SOCIAL RANDOM | Yes |
| | Mental - random interaction | TOM-RANDOM | Yes |
| Emotional Processing | Emotional faces | EMOTION FACES | Yes |
| | Shapes | EMOTION SHAPES | Yes |
| | Emotional faces-shapes | EMOTION FACES-SHAPES | Yes |
| Working Memory | 0-back body | WM 0BK_BODY | Yes |
| | 0-back face | WM 0BK_FACE | Yes |
| | 0-back place | WM 0BK_PLACE | Yes |
| | 0-back tool | WM 0BK_TOOL | Yes |
| | 2-back body | WM 2BK_BODY | Yes |
| | 2-back face | WM 2BK_FACE | Yes |
| | 2-back place | WM 2BK_PLACE | Yes |
| | 2-back tool | WM 2BK_TOOL | Yes |
| | 2-back | WM 2BK | |
| | 0-back | WM 0BK | |
| | 2-back - 0-back | 2BK-0BK | Yes |
| | Body | WM BODY | |
| | Face | WM FACE | |

| | | | |
|---|---|---|---|
| | Place | WM PLACE | |
| | Tool | WM TOOL | |
| | Body - average | WM BODY-AVG | Yes |
| | Face - average | WM FACE-AVG | Yes |
| | Place - average | PLACE-AVG | Yes |
| | Tool - average | TOOL-AVG | Yes |
| Motor | Cue | MOTOR CUE | |
| | Left foot | MOTOR LF | Yes |
| | Left hand | MOTOR LH | Yes |
| | Right foot | MOTOR RF | Yes |
| | Right hand | MOTOR RH | Yes |
| | Tongue | MOTOR T | Yes |
| | Average | MOTOR AVG | |
| | Cue - average | MOTOR CUE-AVG | |
| | Left foot - average | MOTOR LF-AVG | Yes |
| | Left hand - average | MOTOR LH-AVG | Yes |
| | Right foot - average | MOTOR RF-AVG | Yes |
| | Right hand - average | MOTOR RH-AVG | Yes |
| | Tongue - average | MOTOR T-AVG | Yes |
| Gambling | Reward | GAMBLING REWARD | Yes |
| | Punish | GAMBLING PUNISH | Yes |
| | Punish - reward | GAMBLING PUNISH-REWARD | Yes |

**Supplemental Table 2: task contrasts from the Human Connectome Project (HCP).** Details of the imaging and preprocessing protocols are available in Barch et al., 2013. The last column indicates if the task contrasts are also present in the IBC dataset.
